# Escape from SARS-CoV-2 Nsp1-mediated host shutoff by TIAR transcript reveals general features of Nsp1 resistance

**DOI:** 10.1101/2025.08.04.668557

**Authors:** Caleb Galbraith, Madeleine Stolz, Scott Tersteeg, Emily Andrews, Trushar R. Patel, Denys A. Khaperskyy

## Abstract

Severe acute respiratory syndrome coronavirus 2 (SARS-CoV-2) immune escape strategies include general inhibition of host gene expression referred to as host shutoff. Viral non-structural protein 1 (Nsp1) is the main host shutoff factor that blocks protein translation and induces messenger RNA (mRNA) cleavage and degradation. Viral mRNAs are resistant to the translation shutoff and cleavage induced by Nsp1, and the 5’ leader sequence present in all viral mRNAs has been shown to confer resistance. However, the exact molecular mechanism for escape from Nsp1 host shutoff has not been demonstrated. In our previous work, we analyzed the effects of Nsp1 on the expression and function of cellular proteins important for stress granule formation. We discovered that the host transcript for the TIA1 cytotoxic granule-associated RNA binding protein like 1 (TIAL1, commonly referred to as TIAR) is resistant to SARS-CoV-2 Nsp1 host shutoff. In this work, using reporter shutoff assays, we examined sequence and structural features of the TIAR 5’ untranslated region (UTR) and discovered that the first 23 nucleotides of the TIAR transcript are both necessary and sufficient to confer resistance to the Nsp1. Furthermore, our work revealed that the lack of guanosines within a window of 10 to 18 nucleotides downstream from the 5’ end is a defining feature of Nsp1-resistant transcripts shared between the SARS-CoV-2 leader sequence and the TIAR 5’ UTR. Our findings are consistent with the model in which sequence features of 5’ UTRs, rather than their secondary structure, confer resistance to Nsp1 host shutoff to both viral and cellular mRNAs.

## INTRODUCTION

To undermine a host cell’s ability to induce antiviral responses and simultaneously gain preferential access to the host biosynthetic machinery many viruses possess general host shutoff factors (Gaucherand and Gaglia 2022; Bougon and Khaperskyy 2025). Unlike viral inhibitors of specific antiviral sensors, signaling pathways, or effector proteins, viral host shutoff factors target core processes that are required for cellular gene expression, thus broadly undermining a cell’s ability to respond to infection. Host shutoff factors of different RNA and DNA viruses are known to target cellular messenger RNA (mRNA) transcription, processing, nuclear export, or translation (Gaucherand and Gaglia 2022; Bougon and Khaperskyy 2025). Accordingly, shutoff mechanisms must distinguish between host and viral mRNAs to ensure that viral transcription or translation is not affected. In many cases the same mechanisms that ensure that the viral gene expression is not adversely affected allow for select host transcripts to escape host shutoff (Gaucherand et al. 2019; Muller and Glaunsinger 2017; Rao et al. 2021). It is likely that certain host gene products are intentionally spared because they are needed for efficient viral replication (Muller and Glaunsinger 2017; Park et al. 2025). It is also possible that some host transcripts escape viral shutoff due to their inherent properties and have minimal or no effect on the virus.

Severe acute respiratory syndrome coronavirus 2 (SARS-CoV-2), the causative agent of the global coronavirus disease of 2019 (COVID-19) pandemic, encodes several host shutoff factors of which the non-structural protein 1 (Nsp1) is the first to be made in the newly infected cell (Steiner et al. 2024; Schubert et al. 2023; Hartenian et al. 2020). In SARS-CoV-2, like in all Betacoronaviruses, Nsp1 protein is encoded by the first N-terminal portion of the open reading frame 1ab (ORF1ab) that starts being translated directly from the positive-sense viral genomic RNA upon virus uncoating in the cytoplasm and before the onset of replication (Steiner et al. 2024). Betacoronavirus Nsp1 proteins perform their host shutoff function by binding to the ribosome and inhibiting host translation initiation (Schubert et al. 2023). In addition, Nsp1 proteins of SARS-CoV-2 and closely related viruses belonging to subgenus Sarbecovirus (e.g. SARS-CoV and the Middle East respiratory syndrome coronavirus (MERS-CoV)) mediate host mRNA degradation, causing marked depletion of cellular mRNA in infected cells (Kamitani et al. 2006; Lokugamage et al. 2015; Mendez et al. 2021). Structural analyses of SARS-CoV-2 Nsp1 in complex with the small ribosomal subunit revealed in great detail how the Nsp1 C-terminal domain binds and occupies the mRNA entry channel, providing strong evidence for the mechanism of translation inhibition by the Nsp1 (Thoms et al. 2020; Schubert et al. 2020). These structures also suggest that unless the C-terminal domain of the SARS-CoV-2 Nsp1 is displaced from the mRNA entry channel, the ribosome would not be able to translate any mRNA, host or viral. Accordingly, the current model for selective translation of viral transcripts proposes that the conserved structural and/or sequence features of the SARS-CoV-2 mRNA 5’ terminal untranslated region (UTR) specifically interact with the Nsp1 in a manner that results in its displacement from the mRNA entry channel (Mendez et al. 2021; Tidu et al. 2021). While the exact molecular mechanism of the Nsp1-mediated RNA degradation has not been discovered, and it is unclear if a host nuclease is required for the initial Nsp1-mediated target cleavage, the same displacement mechanism that allows for viral mRNA translation could prevent viral mRNA cleavage and degradation (Tardivat et al. 2023; Shehata and Parker 2023).

The positive sense SARS-CoV-2 genome and all subgenomic mRNAs are capped and contain the same 75-nucleotide-long cap-proximal sequence called leader (Steiner et al. 2024; Bujanic et al. 2022). Within the leader sequence, the stem loop 1 (SL1) structure positioned 7 nucleotides from the 5’ end has been determined to be both necessary and sufficient to render reporter transcripts resistant to Nsp1-mediated shutoff (Mendez et al. 2021; Schubert et al. 2020; Bujanic et al. 2022). Therefore, SL1 is proposed to mediate specific leader-Nsp1 interactions essential for Nsp1 displacement from the ribosomal mRNA entry channel. Because of the preferential access to ribosomes inactivated by the Nsp1 and the depletion of competing cellular mRNAs, translation of reporter transcripts containing 5’ leader sequence is not only preserved but is significantly enhanced in the presence of Nsp1 (Aviner et al. 2024).

Several types of host mRNAs have been shown to escape Nsp1-mediated shutoff. Notably, translation of mRNAs containing the terminal oligopyrimidine (TOP) motif is not inhibited by SARS-CoV-2 Nsp1 overexpression (Rao et al. 2021). These mRNAs are characterized by the 5’ terminal cytidine followed by a stretch of pyrimidines in their 5’ UTR (Thoreen et al. 2012). TOP mRNAs preferentially encode ribosomal proteins and translation factors, and their translation is responsive to the activation of the mechanistic target of rapamycin complex 1 (mTORC1) signaling (Thoreen et al. 2012; Jia et al. 2021). This regulation is mediated through mTORC1-mediated phosphorylation of La-related protein 1 (LARP1) that in its unphosphorylated form binds to the 5’ ends of TOP mRNAs and inhibits their translation (Jia et al. 2021; Berman et al. 2021). The exact mechanism of TOP mRNA escape from the Nsp1-mediated shutoff is currently not known.

In our recent work we examined the effects of the SARS-CoV-2 Nsp1-mediated translation inhibition and RNA degradation on the integrated stress response (ISR) and formation of stress granules – biomolecular condensates of mRNA and proteins that form in the cytoplasm following stress-induced inhibition of translation (Kedersha and Anderson 2007; Protter and Parker 2016; Dolliver et al. 2022). Our work has shown that the Nsp1 inhibits both the ISR and stress granule formation (Dolliver et al. 2022). Interestingly, we have also shown that the mRNA degradation by the Nsp1 causes selective depletion of the G3BP stress granule assembly factor 1 (G3BP1) protein, but not other stress granule nucleating proteins. In fact, expression of the stress granule protein TIA1 cytotoxic granule-associated RNA binding protein like 1 (TIAL1, commonly referred to as TIAR) was significantly increased at both the mRNA and protein levels when the Nsp1 was expressed (Dolliver et al. 2022). TIAR is not encoded by a TOP mRNA and does not contain sequences or structure elements that resemble SARS-CoV-2 SL1 and therefore represents a novel type of Nsp1-resistant host transcript. In this work we examined sequence and structural features of the TIAR 5’ UTR and have demonstrated that the first 23 nucleotides (nt) of the UTR are both necessary and sufficient to confer enhanced expression of a reporter in the presence of Nsp1. Furthermore, our work revealed that the lack of guanosines within a window of 10 to 18 nucleotides downstream from the 5’ cap is the defining feature of Nsp1-resistant transcripts shared between the SARS-CoV-2 leader sequence and the TIAR 5’ UTR, but not TOP mRNAs. Using eukaryotic elongation factor 2 (EEF2) 5’UTR reporter as a model TOP mRNA we have shown that despite the lack of the G-less window 10 to 18 nucleotides from the cap, this reporter escapes Nsp1-mediated host shutoff, and this escape does not require LARP1. Altogether, our work reveals two independent mechanisms of resistance from Nsp1-mediated host shutoff and provides an alternative model for viral mRNA escape that is not dependent on SL1 structure.

## RESULTS

### TIAR 5’ UTR is sufficient for escape from Nsp1 host shutoff

The Nsp1 resistance determinants of the viral transcripts and the host TOP mRNAs are located in their 5’ UTRs. Therefore, we started our analyses of the putative resistance determinants in the TIAR mRNA by inserting its 499-nucleotide-long 5’ UTR upstream of the mScarlet reporter construct. Then we compared reporter expression in HEK 293A cells co-transfected with the Nsp1 expression construct or the control luciferase encoding vector. Reporters containing the SARS-CoV-2 (CoV2) 5’UTR and the control vector-derived 5’ UTR upstream of the mScarlet coding sequence, previously characterized by Vora et al. (Vora et al. 2022), served as the Nsp1 resistant and the Nsp1 sensitive benchmarks (Figure 1A). As expected, Nsp1 co-expression dramatically reduced mScarlet signal from the control 5’ UTR reporter. By contrast, both the CoV2 and the TIAR 5’ UTRs increased mScarlet expression in the Nsp1 co-expressing cells compared to vector control (Figure 1B-D). Notably, the basal expression of these Nsp1-resistant reporters was significantly reduced compared to the control, but in the presence of Nsp1, it reached levels comparable to the basal level of the control 5’ UTR reporter (Figure 1C).

**Figure 1.**
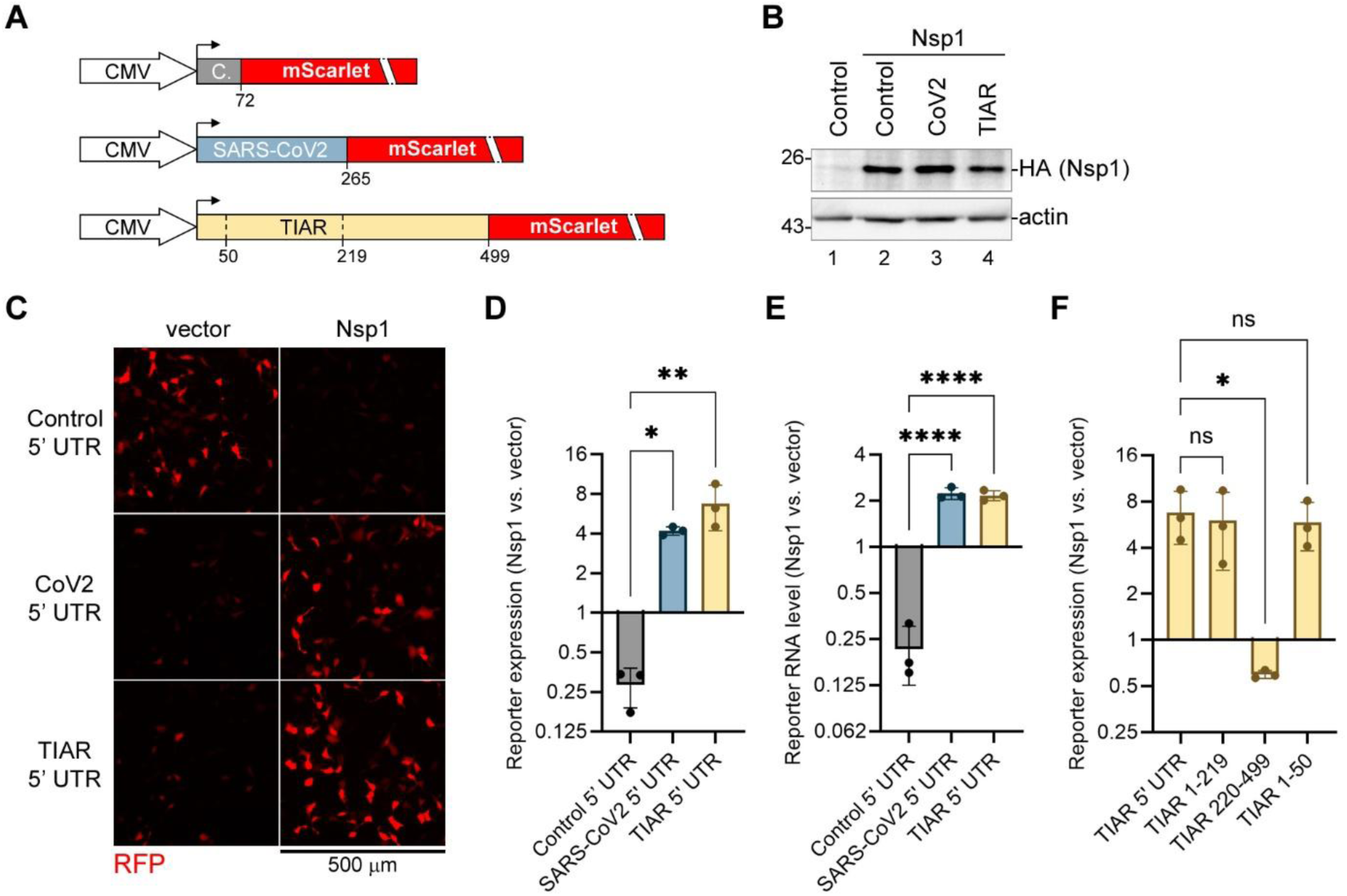
First 50 nucleotides of TIAR transcript are responsible for Nsp1-mediated increase in expression. (A) Schematic diagram of reporter constructs used in this study showing the CMV promoter followed by the indicated 5’ UTR sequences upstream of the mScarlet open reading frame (C.= Control UTR). Numbers indicate UTR lengths and positions that flank TIAR 5’ UTR deletions tested in this work. (B-E) 293A cells were co-transfected with reporters shown in panel A and the expression construct for HA-tagged Nsp1 or the control vector. (B) Expression of the Nsp1 protein when co-transfected with the indicated 5’ UTR reporter constructs was analysed by western blot using anti-HA antibody. (C) At 24 h post transfection, reporter expression was analysed by fluorescence microscopy. Scale bar = 500 µm. (D) The relative signal intensity in the RFP channel was quantified from images represented in panel B to measure down- or upregulation by Nsp1 compared to the control vector (N = 3, see materials and methods section for details). (E) Relative levels of the indicated mScarlet reporter RNAs were analyzed using RT-qPCR (N = 3). (F) Relative expression of the indicated TIAR 5’ UTR mScarlet reporters was quantified (N = 3). In D,E and F, one-way ANOVA and Dunnett multiple comparisons tests were used to determine statistical significance (ns: non-significant; ***P-value <0.001; **P-value < 0.01; *P-value < 0.05). Error bars = standard deviation.

In our previous work, we have shown that the CoV2 Nsp1 expression increases the endogenous TIAR mRNA levels (Dolliver et al. 2022). Therefore, we determined the relative levels of our three UTR reporter transcripts in the Nsp1 and the vector control expressing cells using reverse transcription quantitative PCR (RT-qPCR). Our analyses showed that the CoV2 and the TIAR 5’ UTR reporter transcript levels were significantly increased by the Nsp1, while the Nsp1 sharply reduced the control reporter mRNA levels (Figure 1E). Thus, the 5’ UTR of TIAR was sufficient to confer the Nsp1 resistance to a heterologous reporter and to mediate the increased translation and increased mRNA levels in the presence of the Nsp1 similar to the endogenous TIAR transcript.

Next we wanted to determine which region of the TIAR 5’ UTR is responsible for conferring Nsp1 resistance by constructing reporters with large deletions of the 5’ terminal and the 3’ terminal portions of the sequence. Nsp1 co-expression experiments with these reporters showed that the first 50 nt of the TIAR 5’ UTR (TIAR 1-50) are sufficient for the Nsp1 protection and enhanced mScarlet expression, comparable to that of the full 499-nt 5’ UTR (Figure 1F).

### Stem loops found in TIAR 5’UTR are dispensable for protection

Since the first 50 nt of the 5’ UTR were functionally equivalent to the full-length TIAR 5’ UTR in conferring increased expression in the presence of Nsp1, we focused our subsequent analyses on this sequence. First, we compared the predicted secondary structure of TIAR 1-50 to the corresponding portions of the CoV2 5’ UTR that contains SL1 and the Control vector UTR (Figure 2A). The predicted RNAfold minimal free energy structure showed two stem loops in the TIAR 1-50 sequence: a small stem loop at the very 5’ end that we termed SL1 and a longer stem loop (SL2) starting 23 nt downstream from the 5’ end. Overall secondary structures of the three 5’ UTRs looked distinct, with either SL1 or SL2 in TIAR 1-50 lacking similarity to SL1 of CoV2. Interestingly, a polypyrimidine sequence 5’-CUUCCC-3’ found at the tip of CoV2 SL1, which was shown to be essential for its function in mediating Nsp1 resistance (Bujanic et al. 2022), is also present in TIAR 1-50; however, it is not a part of a stem loop (Figure 2A). Because the current model suggests that specific binding of the Nsp1 N-terminal domain to the CoV2 SL1 is required for shutoff evasion by the viral transcripts, we compared the binding affinities of the purified full-length Nsp1 or just its 127 amino acid long N-terminal domain (N127) to the in vitro transcribed model RNAs corresponding to the first 50 nt of the three 5’ UTR sequences using microscale thermophoresis (Figure 2B-C). Our analyses showed that the full-length Nsp1 had higher affinity for the model RNAs compared to N127 (Figure 2C). Importantly, Nsp1 bound with higher affinity to CoV2 and TIAR 5’ UTR RNAs compared to Control UTR RNA, and for each of the Nsp1 proteins, the binding was similar for the TIAR sequence compared to CoV2 (K_d_ = 4.23 µM vs. 6.30 µM for the full-length Nsp1 and 112.4 µM vs. 77.83 µM for the N127, respectively). Thus, despite having distinct sequence and the secondary structure, both the first 50 nucleotides of the TIAR and CoV2 5’ UTR were able to bind Nsp1 in the absence of any other cellular factors, demonstrating that the CoV2 SL1 does not uniquely confer higher affinity binding under these conditions.

**Figure 2.**
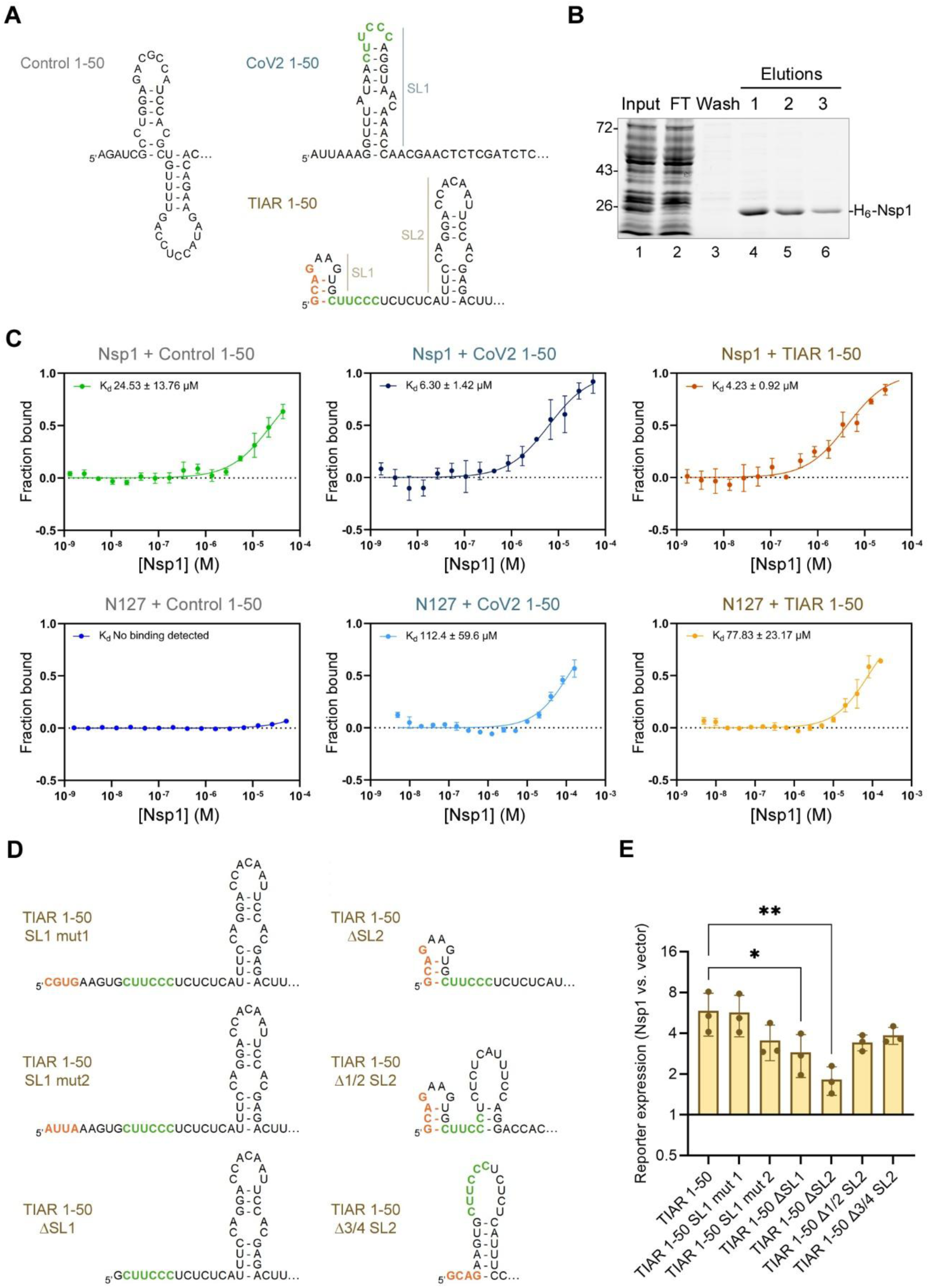
Stem loop structure of the TIAR 5’ UTR is not important in protection from Nsp1 shutoff. (A) Comparison of predicted secondary structures of the first 50 nt of the CoV2 and TIAR RNAs compared to the Control vector 5’ UTR. Structures are drawn based on RNAfold minimum free energy predictions (see materials and methods section for details). Identical 6-nt pyrimidine stretches are colored green and the 5’ terminal nucleotides forming part of the stem loop 1 (SL1) of TIAR are colored orange for reference. SL2 = stem loop 2. (B) Input, flow through (FT), first wash, and elution fractions from His-tagged Nsp1 protein purification were separated and visualised on a 10% Stain Free polyacrylamide gel. (C) Microscale thermophoresis assay binding curves for the purified full length (top) and the C-terminally truncated (N127, bottom) Nsp1 proteins and synthetic RNAs corresponding to the first 50 nt of the Control, CoV2 and TIAR transcripts. (D) Predicted secondary structures of the indicated TIAR 5’ UTR modifications tested in reporter assays. Structures are drawn based on RNAfold minimum free energy predictions (see materials and methods section for details). Coloring of specific regions corresponds to panel A. (E) 293A cells were co-transfected with the indicated TIAR 5’ UTR mScarlet reporters and the expression construct for HA-tagged Nsp1 or the control vector. Relative reporter expression was quantified and plotted (N = 3), one-way ANOVA and Dunnett multiple comparisons tests were used to determine statistical significance (**P-value < 0.01; *P-value < 0.05). Error bars = standard deviation.

Next, we constructed a series of TIAR 1-50 deletion and substitution mutants and tested the corresponding mScarlet reporter constructs in our co-transfection shutoff assay to determine which secondary structure features were important for escape from the Nsp1 shutoff (Figure 2D-E). First, we disrupted TIAR SL1 by replacing the first four nucleotides with random ones (SL1 mut1) or with the first 4 nucleotides of the CoV2 leader (SL1 mut2). RNAfold prediction confirmed the lack of SL1 formation in both mutants. Then, we performed deletions of the entire SL1, entire SL2, half of SL2, or ¾ of SL2 sequences (Figure 2D). Remarkably, all of these mutants’ 5’ UTR constructs were resistant to the Nsp1-mediated shutoff (Figure 2D). This suggests that the sequence features 9-23 nt from the 5’ end of TIAR 1-50 (before beginning of SL2), rather than the secondary structures of either SL1 or SL2 are responsible for protection.

### A G-less window in the 5’ UTR in both TIAR and SARS-CoV-2 leader-containing transcripts mediates Nsp1 resistance

Sequences 9-23 nt from the 5’ end in both the CoV2 and TIAR transcripts contain the identical 6-pyrimidine stretch 5’-CUUCCC-3’. To test if this polypyrimidine sequence is necessary for conferring resistance to the Nsp1-mediated shutoff, we deleted the pyrimidine stretch from the TIAR 1-50 5’ UTR reporter (producing TIAR 1-50 Δ12-21 reporter) or the entire first 23 nt from the TIAR 1-50 (producing TIAR 23-50 reporter). We also created reporters with C to G substitutions at positions 14 and 19 of the TIAR 1-50 that interrupt its 12-nt pyrimidine stretch, including the sequence also present in the CoV2 leader. To our surprise, only the last mutation that introduced guanosines to disrupt the pyrimidine stretch rendered the reporter construct sensitive to the Nsp1-mediated shutoff (Figure 3A). The expression of the deletion mutant reporters was no longer significantly increased by the Nsp1, but they remained completely resistant to the Nsp1 shutoff. To verify that the introduction of guanosines within the pyrimidine tract is sufficient to sensitize the full-length TIAR 5’ UTR reporter to Nsp1, we constructed the full-length TIAR 5’ UTR C14,19G mutant reporter and observed that it was similarly downregulated by the Nsp1 (Figure 3A). When we aligned different sequences of all reporters that we tested so far in our mScarlet shutoff assay, we saw that all Nsp1-resistant constructs had a G-less window positioned 10-18 nt from the 5’ end, while all sensitive reporters had guanosines within that window (Figure 3B). Indeed, our pyrimidine stretch deletions resulted in repositioning of adenosines but not guanosines within 10 to 18 nt from the 5’ end. Altogether, our results strongly suggest that the lack of guanosines 10 to 18 nt downstream from the 5’ cap is both necessary and sufficient to protect TIAR-derived 5’ UTR reporter from the Nsp1-mediated shutoff.

**Figure 3.**
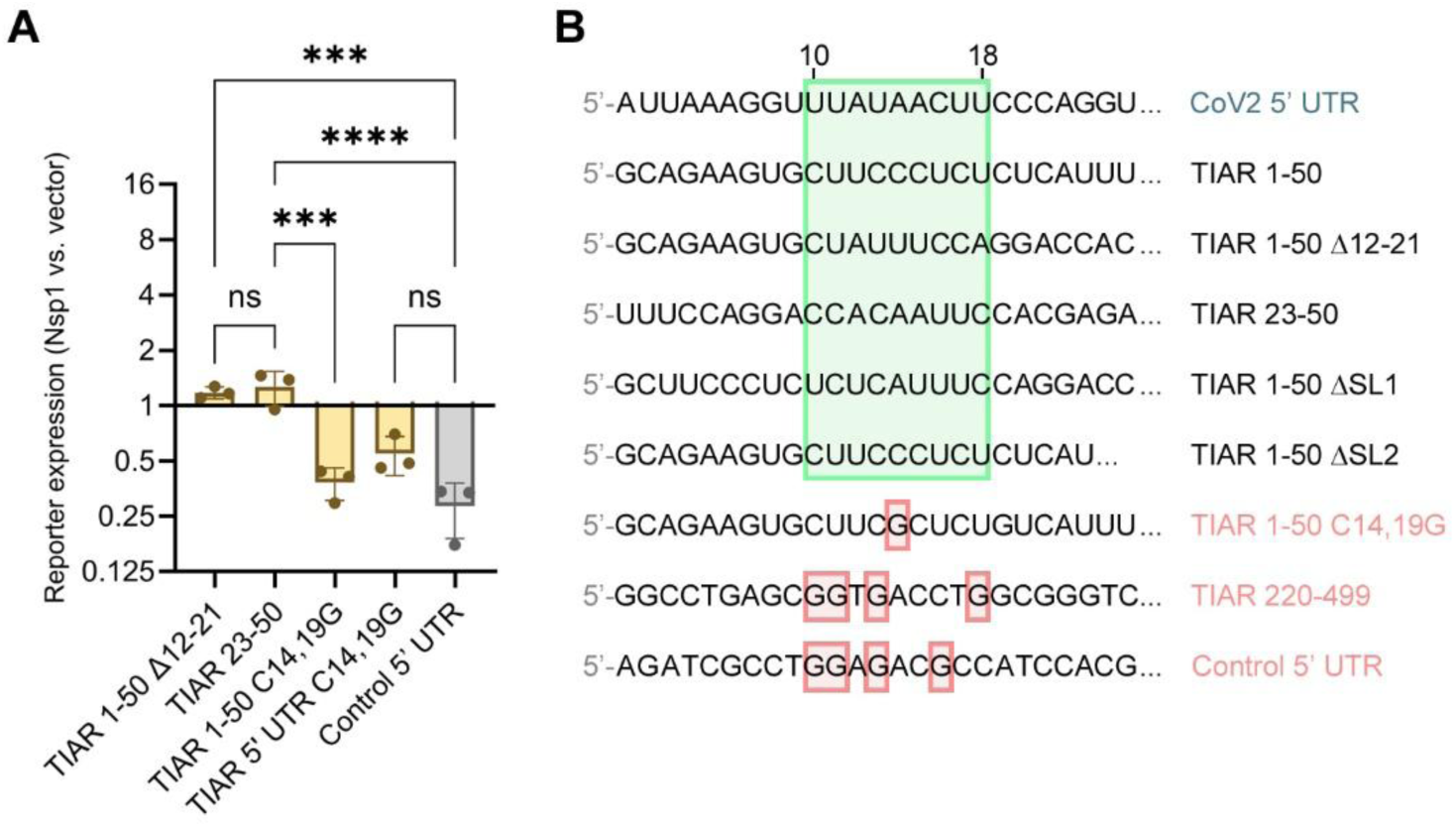
Guanosines within a defined window downstream of the 5’ cap determine sensitivity to Nsp1 shutoff. (A) 293A cells were co-transfected with the mScarlet reporters with the indicated 5’ UTRs and the expression construct for HA-tagged Nsp1 or the control vector. Relative reporter expression was quantified and plotted (N = 3), one-way ANOVA and Tukey multiple comparisons tests were used to determine statistical significance (ns: non-significant; ***P-value < 0.001; ****P-value < 0.0001). Error bars = standard deviation. (B) Cap-proximal sequences of the indicated 5’ UTRs that confer resistance (CoV2 and TIAR-derived sequences labeled with black text) or sensitivity (labeled with red text) to Nsp1. A G-less window positioned 10 to 18 nucleotides from the transcription start site in all Nsp1-resistant UTRs is indicated with a green box. Guanosines within the corresponding window of the Nsp1-sensitive UTRs are highlighted with red boxes.

### Terminal oligopyrimidine motif reporter mRNA has a distinct LARP1-independent mechanism of Nsp1 resistance

Another type of Nsp1-resistant transcripts is the TOP mRNAs, and several TOP 5’ UTRs tested previously in Nsp1 reporter shutoff assays were demonstrated to be sufficient to mediate Nsp1 resistance (Rao et al. 2021). However, only a 5 pyrimidine stretch immediately following the invariant initiating 5’ cytosine is necessary for the transcript to be defined as TOP mRNA. To test the TOP mRNA reporter in our assay, we cloned the first 50 nt of the eukaryotic elongation factor 2 (EEF2) 5’ UTR upstream of the mScarlet coding sequence. This 5’ UTR contains 3 evenly spaced guanosines within the window of 10-18 nt downstream from the 5’ cap which distinguishes this sequence from our set of the Nsp1-resistant 5’ UTRs (Figure 4A). Nevertheless, consistent with previous studies, the EEF2 5’ UTR reporter was also resistant to the Nsp1 shutoff in our system (Figure 4B-C). TOP mRNA 5’ ends can be bound by the LARP1 protein (Berman et al. 2021); therefore, one possible mechanism for their resistance to Nsp1 could be that LARP1 interferes with Nsp1 shutoff function. To test this hypothesis, we compared the effects of Nsp1 on the EEF2 5’ UTR reporter in 293A cells in which LARP1 expression was silenced using two different shRNA constructs or the control cells expressing shRNA targeting interleukin 4 (IL4), which is not expressed in these cells. We also included CoV2 and TIAR 1-50 5’ UTR reporters in these assays. First, we confirmed successful LARP1 knock down using western blotting (Figure 4D). Both shRNA constructs downregulated LARP1 protein levels, with the second shRNA (LARP1-2) having a stronger effect (Figure 4D, lanes 5,6). Then, we analyzed and quantified the mScarlet reporter expression and did not observe significant effects of the LARP1 silencing on any of the Nsp1-resistant reporters (Figure 4E). Subsequently, we performed 5’ rapid amplification of cDNA ends (5’RACE) analysis and confirmed that transcription of our EEF2 5’ UTR reporter initiates at the canonical TOP 5’ cytosine (Figure 4F). Together, these results reveal that the LARP1 is not required for the TOP mRNA escape from shutoff by the CoV2 Nsp1; therefore, a distinct mechanism must exist for protection of TOP mRNAs from the Nsp1 that permits the presence of guanosines 10-18 nt from the 5’ end of the transcript.

**Figure 4.**
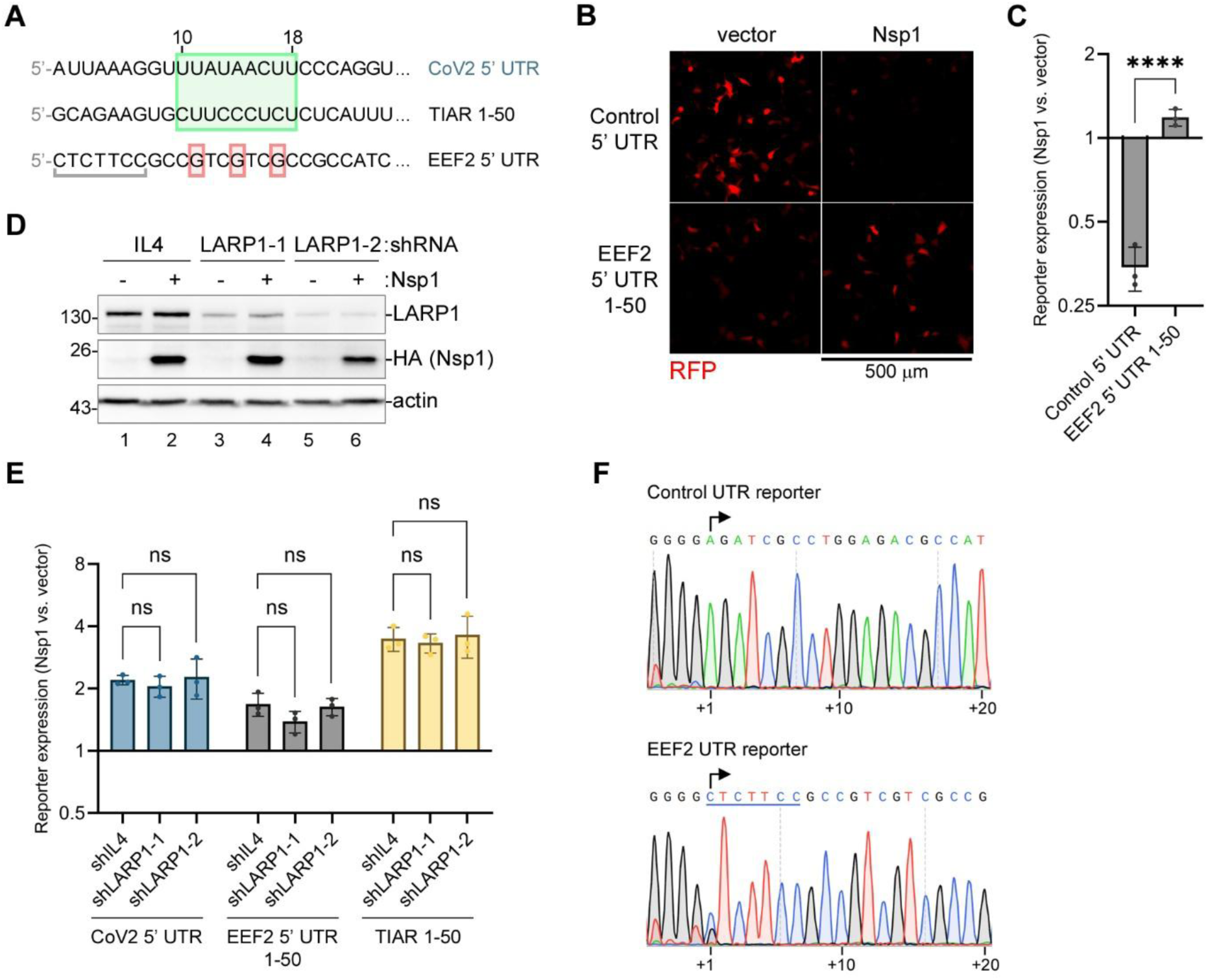
Protection from Nsp1 shutoff is LARP1-independent. (A) Cap-proximal sequences of the indicated 5’ UTRs that confer resistance (CoV2, TIAR, and EEF2) to Nsp1. A G-less window positioned 10 to 18 nucleotides from the transcription start site in CoV2 and TIAR UTRs is indicated with a green box. Guanosines within the corresponding window of the EEF2 transcript are highlighted with red boxes. Terminal oligopyrimidine motif in EEF2 is underlined. (B,C) 293A cells were co-transfected with the indicated reporters and the expression construct for HA-tagged Nsp1 or the control vector. (B) At 24 h post transfection, reporter expression was analysed by fluorescence microscopy. Scale bar = 500 µm. (C) The relative signal intensity in the RFP channel was quantified from images represented in panel B to measure down- or upregulation by Nsp1 compared to the control vector (N = 3, see materials and methods section for details). (D,E) 293A cells expressing shRNAs targeting LARP1 or control shRNA targeting IL4 were co-transfected with the mScarlet reporters and the expression construct for HA-tagged Nsp1 or the control vector. (D) Levels of LARP1 and HA-Nsp1 proteins was analysed by western blot at 24 h post transfection. (E) Relative expression of the indicated reporters was quantified and plotted (N = 3), one-way ANOVA and Dunnett multiple comparisons tests were used to determine statistical significance (ns: non-significant). Error bars = standard deviation. (F) Transcription start sites for the EEF2 and control reporters were determines using 5’ RACE. Sanger sequencing traces downstream of the guanosine stretch that corresponds to oligo(dC) tail added to the cDNA end by the reverse transcriptase are shown. Arrows indicate the mapped transcription start site positions. Terminal oligopyrimidine sequence in EEF2 UTR reporter is underlined.

### Sensitivity to Nsp1 shutoff does not correlate with basal reporter expression level

Throughout our analyses of various reporter constructs we often observed varied but generally lower basal expression of the Nsp1-resistant constructs in the absence of the Nsp1 compared to our Nsp1-sensitive control reporter (Figures 1C, 4B). In the presence of Nsp1, their expression increased in many but not all cases. This led us to investigate whether increased expression of the Nsp1-resistant reporters results mainly from alleviated competition from efficiently translated host mRNAs depleted by the Nsp1, and whether the degree of upregulation inversely correlates with the basal expression level. For this analysis, we compiled raw mScarlet fluorescence measurement data from all our experiments and normalized expression in the absence of the Nsp1 to our control reporter levels. Then, we plotted these values versus fold change due to Nsp1 co-expression (Figure 5). Indeed, we observed that our control reporter had the highest basal expression levels and was the most potently downregulated by the Nsp1. The CoV2 5’ UTR reporter, by comparison, had on average 4 times lower basal expression and was upregulated on average 4-fold by the Nsp1. However, the TIAR 5’ UTR reporter had higher basal expression and was even more strongly upregulated by Nsp1 co-expression (approximately 6-fold) than the CoV2 5’ UTR reporter. Overall, we did not observe a significant correlation between the basal expression level and the Nsp1-mediated upregulation or downregulation in our dataset (Pearson r = −0.207, p value 0.425), with some of the mutant TIAR 5’ UTR reporters having similar basal expression levels but the opposite response to the Nsp1 (Figure 5). It is still possible that the same 5’ UTR features that cause highly efficient translation initiation and/or increased mRNA stability could confer sensitivity to the Nsp1-mediated shutoff; however, the opposite is not true for the specific transcripts that are more efficiently translated in the presence of the Nsp1.

**Figure 5.**
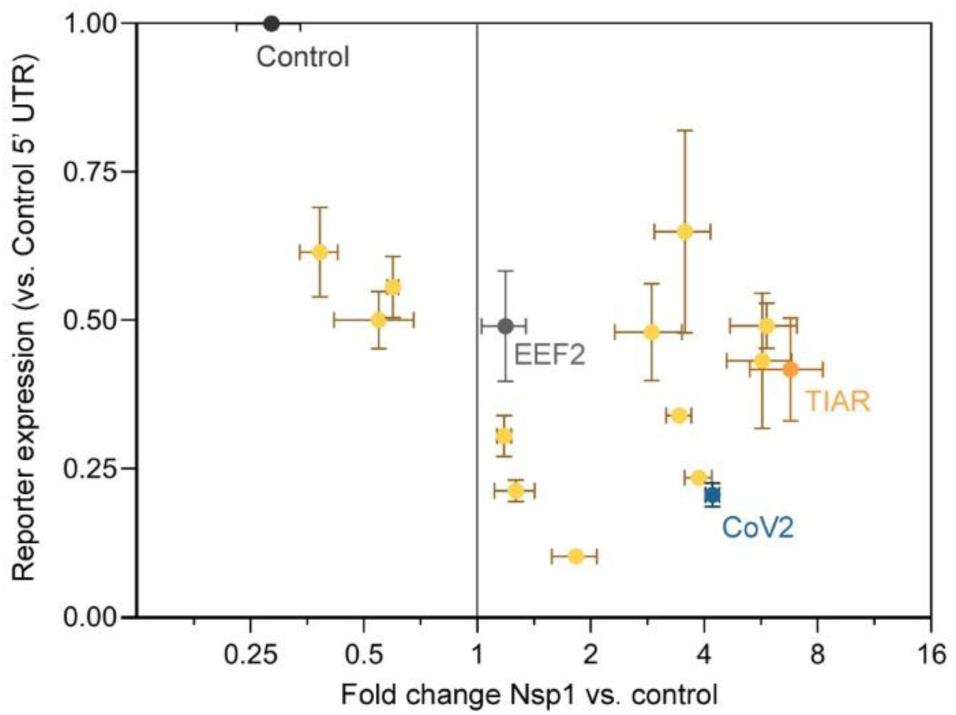
Lack of Nsp1-mediated shutoff does not correlate with low basal level of reporter expression. Basal 5’ UTR reporter expression levels (in the absence of Nsp1) measured in this study were normalized to the basal Control 5’ UTR reporter expression and plotted versus their Nsp1 coexpression-induced fold change. Different mutant TIAR 5’ UTR reporters analysed in figures 1-3 (orange dots) are not labeled individually. Error bars = standard deviation.

## DISCUSSION

Despite significant continued research effort towards understanding the mechanisms that allow SARS-CoV-2 to successfully subvert host innate and adaptive immunity, including host shutoff, the molecular mechanisms for the Nsp1-mediated translation inhibition and mRNA degradation remain poorly characterized. Specifically, the exact mechanism of the mRNA cleavage induced by the Nsp1 has not been established and whether the Nsp1 can itself act as a nuclease remains uncertain, despite recent studies suggesting that under certain conditions it can cleave RNA (Tardivat et al. 2023; Abaeva et al. 2023). Similarly, ample evidence indicates that the 5’ leader sequence present in all viral mRNAs is responsible for rendering these mRNAs resistant to shutoff by the Nsp1, with SL1 structure and sequence playing the major role (Mendez et al. 2021; Bujanic et al. 2022; Vora et al. 2022). However, the exact molecular mechanism for escape from the Nsp1 shutoff has not been demonstrated. In our previous work we serendipitously discovered that the host transcript for the stress granule protein TIAR is resistant to the SARS-CoV-2 Nsp1 shutoff and the TIAR protein expression is increased in the Nsp1-expressing cells compared to control cells (Dolliver et al. 2022). Importantly, we did not observe increased TIAR protein or mRNA levels in SARS-CoV-2 infected cells, indicating that it is not specifically upregulated by the virus and is subject to shutoff by viral factors other than Nsp1 (e.g. Nsp16 (Banerjee et al. 2020), ORF6 (Addetia et al. 2021)). Because TIAR mRNA 5’ UTR lacked any known Nsp1 resistance elements, in the present work we set out to analyze its structural and sequence features that mediate Nsp1 resistance using a series of reporter assays. Our minimal system is well suited for comparison of different 5’ UTRs because other parts of the reporter (promoter, coding sequence, 3’ UTR) remain constant and our control construct is strongly inhibited by the Nsp1. Our experiments have shown that the first 23 nucleotides of the TIAR 5’ UTR are sufficient to confer Nsp1 resistance, and that the G-less stretch 10 to 18 nt from the 5’ end is necessary for this effect. By contrast, the two stem loop structures predicted to form in the first 50 nt of the TIAR 5’ UTR did not have any measurable effect. Therefore, our results suggest that the positioning of sequences relative to the 5’ cap strongly influences susceptibility to Nsp1, rather than the presence of specific secondary structures. Our data is in agreement with the study by Bujanic et al. (Bujanic et al. 2022), in which they introduced substitutions in the SL1 sequence of the CoV2 5’ UTR and observed that the introduction of adenosines, cytidines, or uridines that disrupted structural features did not abolish the Nsp1 resistance of reporter constructs. By contrast, introduction of guanosines that either disrupted or preserved the SL1 architecture was sufficient to render reporters susceptible to Nsp1 shutoff (Bujanic et al. 2022). A more recent study by Berlanga et al. (Berlanga et al. 2025) that was published when our manuscript was in preparation has shown that an entirely G-less 5’ UTR confers both the Nsp1 resistance and enhanced expression of a reporter, confirming our main finding that the guanosines in the 5’ UTR make the transcript sensitive to Nsp1 shutoff. Presence of guanosines in 5’ UTR increases formation and stability of secondary structures that have to be unwound by the helicase activity of the eukaryotic translation initiation factor 4A (eIF4A), a component of the eIF4F initiation complex (Brito Querido et al. 2024a). It is possible that the eIF4A activity is required for the Nsp1 shutoff mechanism. As an integral part of the eIF4F complex, eIF4A is recruited into the 48S preinitiation complex even in the absence of strong RNA secondary structures, however a recent structural analysis of the 48S preinitiation complex reveals a second molecule of eIF4A away from the eIF4F that interacts with the ribosome near the RNA entry channel, which is in close proximity to the site occupied by the Nsp1 (Brito Querido et al. 2024b). This is consistent with the helicase action of the eIF4A that would be required at the RNA entry to the ribosome rather than at the exit site where eIF4F is positioned. Notably, the second eIF4A in that structure was in close proximity to the eIF3G RNA recognition motif that can serve as a co-factor in the RNA cleavage by the purified Nsp1 (Abaeva et al. 2023). Another possibility consistent with the sequence rather than the secondary structure being crucial for the mRNA sensitivity to the Nsp1 is that the G-less sequences are more efficient at outcompeting the Nsp1 C-terminal domain in the RNA entry channel of the ribosome. This could be due to interactions with the Nsp1 itself or with a yet to be identified RNA binding protein. In our work, binding of both the TIAR and CoV2 5’ UTR RNAs in the absence of the ribosome or other host proteins was stronger to the full-length protein compared to just the Nsp1 N-terminal domain. This may indicate that the C-terminal domain interactions are important for the Nsp1 displacement from and preferential translation of resistant transcripts. These analyses together with our reporter co-expression experiments revealed that the TIAR 5’ UTR and CoV2 5’ UTR are functionally equivalent towards the Nsp1 despite having no structural homology and only a 6-nt pyrimidine stretch in common. This pyrimidine stretch, however, was dispensable for the protection from the Nsp1 as long as the G-less window 10 to 18 nt from the 5’ end was present. By contrast, the EEF2 5’ UTR reporter representing a model TOP mRNA had 3 guanosines within this window yet remained resistant to the Nsp1 shutoff. This feature was not dependent on LARP1 binding to the TOP element, as silencing of LARP1 had no effect on reporter expression in the presence of Nsp1. It is possible that the TOP element alters the way the mRNA is presented to the preinitiation complex, leading to Nsp1 displacement through a distinct mechanism.

By using a simple reporter shutoff assay based on ectopic Nsp1 co-expression we were able to focus on the basic 5’ UTR sequence features of the Nsp1 resistant TIAR mRNA. This allowed us to identify a simple sequence feature sufficient for rendering this transcript resistant to the Nsp1 – a G-less sequence 10-18 nt downstream from the 5’ end. However, the analysis of the EEF2 5’ UTR suggests that while sufficient, this G-less window is not the only determinant of protection. We recognise that our study was conducted in a single cell line model, which represents a significant limitation. Furthermore, reliance on a single reporter system, with all its advantages, is another limitation of our study. At the same time, our main findings are consistent with the recent reports by other groups and provide further evidence that the G-less sequences in the 5’ UTR, rather than defined stem loop structures, confer resistance to Nsp1 host shutoff to viral and host mRNAs.

## MATERIALS AND METHODS

### Cell lines

Human embryonic kidney (HEK) 293A and 293T cells were purchased from American Type Culture Collection (ATCC, Manassas, VA, USA) and cultured in Dulbecco’s modified Eagle’s medium (DMEM) supplemented with heat inactivated 10% fetal bovine serum (FBS) and 2 mM L-glutamine (all purchased from Thermo Fisher Scientific (Thermo), Waltham, MA, USA). For LARP1 silencing, shRNA inserts targeting the human LARP1 gene (shLARP1-1, target sequence gcgccagattgaatactactt; shLARP1-2, target sequence gcaagaatacctcggcaaatt) or control human interleukin 4 (IL4) that is not expressed in epithelial cells (shIL4, target sequence agctgatccgattcctgaaac) were cloned into pLKO.1-TRC vector (Addgene plasmid #10878, a gift from David Root, (Moffat et al. 2006)). Then, 293A cells were transduced with lentiviruses generated with the above vectors at an MOI of 1.0, and stably transduced cells were selected with 1 μg/mL puromycin for 48 h. Resistant cells were seeded onto 12-well cluster dishes and used in experiments the following day.

### Plasmids

Generation of an N-terminally HA-tagged SARS-CoV-2 Nsp1 expression vector (pCDNA3-HA-Nsp1) is described in (Dolliver et al. 2022). The pCMV-LUC2CP/ARE control vector expressing firefly luciferase is a kind gift from Gideon Dreyfuss (Addgene plasmid #62857 (Younis et al. 2010)). The mScarlet reporter plasmids with vector-derived control UTR and SARS-CoV-2 (CoV2) 5’ UTRs were constructed from pLV-mScarlet-Control-UTR and pLV-mScarlet-CoV2-UTR (a kind gift from Hao Wu, Addgene plasmids #184638 and #184637, (Vora et al. 2022)) by transferring sequences between the NdeI and EcoRI sites into a pCR3.1-EGFP vector (Khaperskyy et al. 2016) to generate pCR3-UTR-CoV2-mScarlet and pCR3-UTR-Control-mScarlet plasmids. Subsequently, to facilitate the precise insertion of other 5’ UTRs, the pCR3-UTR-Esp-mScarlet recipient vector was generated by simultaneous removal of the CoV2 5’ UTR from the pCR3-UTR-CoV2-mScarlet and insertion of two adjacent Esp3I restriction sites immediately downstream of the transcription start site via PCR mutagenesis. Primers used in the generation of this vector: UTR-Esp-For 5’-tacgtctccatggtgagcaagggcgag and UTR-Esp-Rev 5’-gacgtctcaggttcactaaacgagctctgc. To generate the pCR3-UTR-TIAR-mScarlet and pCR3-UTR-EEF2-mScarlet vectors, synthetic DNA fragments encoding full-length TIAR or the first 50 nt of the EEF2 5’ UTR flanked by Esp3I restriction sites were ordered from Invitrogen (Thermo, Waltham, MA) and cloned into the pCR3-UTR-Esp-mScarlet. TIAR 5’ UTR deletion and substitution mutants were generated by PCR mutagenesis of the pCR3-UTR-TIAR-mScarlet vector. All wild-type and mutant TIAR 5’ UTR sequences are presented in Figures 2-4. pET28-6His-Nsp1 *E.coli* expression vector was generated by inserting the Nsp1 open reading frame from the pcDNA3-HA-Nsp1 into the pET28a+ vector (EMD Biosciences) using EcoRI and XhoI restriction sites and subsequent removal of the thrombin cleavage site sequence using PCR mutagenesis. pET28-6His-Nsp1(N127) was generated from pET28-6His-Nsp1 by deletion of the Nsp1 linker and C-terminal domain coding sequences using PCR mutagenesis. Full sequences of mutagenesis primers and vectors featured in this work are available upon request.

### Minimum free energy UTR structure prediction

RNAfold (R et al. 2011) was used to predict the secondary structure of RNA sequences. To confirm the accuracy of the program, RNA sequences with solved secondary structures (e.g. SARS-CoV-2 5’ UTR) were modelled. Structures were predicted for RNA sequences of the first 50 bases or less in each UTR with default parameters (avoid isolated base pairs, dangling energies on both sides of a helix in any case, 2004 Turner model RNA parameters).

### Reporter transfection

HEK 293A cells were seeded into 20-mm wells of 12-well cluster dishes with or without glass coverslips and transfected the next day with 500 ng DNA mixes/well containing 100 ng reporter vectors, 150 ng pUC19 filler DNA, 250 ng of either the N-terminally HA-tagged Nsp1 expression plasmid pcDNA3-HA-Nsp1 or control firefly luciferase expression plasmid pCMV-LUC2CP/ARE, and 1.5 ug of linear polyethylenimine (Cedarlane, Burlington, ON, Canada) in 100 ul of Opti-MEM-I media (Thermo). Cells were analysed 23–24 h post-transfection as indicated.

### Fluorescence microscopy and mScarlet reporter expression measurement

At 24 h post-transfection, live cells were imaged in the RFP channel using EVOS XL Core Imaging System (Thermo). For each independent biological replicate, duplicate wells were transfected with the same reporter DNA mix, and at least 2 separate random fields of view were captured per well. Total red fluorescence signal was measured for each field of view using ImageJ Fiji software after automatic background subtraction (Schindelin et al. 2012). The integrated density of all pixels in the total of at least four images per replicate were averaged and used as a raw reporter expression value in further analysis. At least 3 independent biological replicates were performed per condition and the normalized values for each replicate were displayed on bar graphs.

### Nsp1 protein purification

BL21 (DE3) *E. coli* cells (Thermo) were transformed with pET28-6His-Nsp1 or pET28-6His-Nsp1(N127) plasmids and maintained in LB media (BioShop) supplemented with 50 μg/mL kanamycin (BioShop). Liquid cultures for protein production were grown to an optical density (OD600) of 0.5-1.2, followed by induction with 1 mM isopropylthio-β-galactoside (IPTG, BioShop) at +20°C overnight. Harvested cell pellets were resuspended in phosphate-buffered saline (PBS) pH7.4 with 10 mM imidazole and 1x EDTA-free protease inhibitor cocktail (all Millipore Sigma) and lysed by two cycles of liquid nitrogen freeze-thawing, followed by four 10-second intervals of sonication. Debris was spun down at 4,000 x g for 20 min and supernatants were combined with HisPur Ni-NTA resin (Fisher Scientific) at 4°C for 30 min, washed three times with 25 mM imidazole in PBS, and eluted three times with 250 mM imidazole in PBS. Presence of the recombinant protein in the elution fractions was confirmed by polyacrylamide gel electrophoresis (PAGE) followed by visualization with Stain Free reagent (Bio-Rad). Elution fractions (3 ml total) were combined and the H6-Nsp1 proteins were further purified by size exclusion chromatography using an ÄKTA Pure FPLC System (Cytiva) on a HiLoad 16/600 Superdex 75 column (Amersham Biosciences), followed by concentration using Amicon Ultra Centrifugal Filter Units (EMD Millipore) with a molecular weight cutoff of 10,000 Daltons. Protein concentration was measured by the DC protein assay (Bio-Rad).

### RNA synthesis and labeling

Template sequences for the SARS-CoV-2 and TIAR 5’ 50-nt model RNAs were inserted in a pUC-GW-Kan plasmid (Gemmill et al. 2024) flanked by a T7 RNA polymerase promoter on the 5’ end and an XbaI restriction endonuclease cut-site sequence on the 3’ end. The template for the vector-derived control 50-nt RNA was PCR-amplified from pCR3-UTR-Control-mScarlet vector using the forward primer that includes a T7 promoter (5’-taatacgactcactatagagatcgcctggagac-3’) and a reverse primer (5’-gtgtcttctatggaggtc-3’) using 2X Taq Master Mix (NEB). The three templates were used in an *in vitro* transcription reaction using in-house purified T7 RNA polymerase for 24 hours at +37°C, followed by a one-hour DNase I digestion. Post digestion, the RNA was ethanol precipitated, resuspended in nuclease-free water (Thermo) and purified using an RNA clean-up kit (NEB). Final RNA elution was in RNA buffer (20mM HEPES pH 6.8, 5mM MgCl2, 150mM NaCl). Purified RNA was subjected to 5’ labelling with the fluorophore Alexa 488 (Thermo). Approximately 7.5 μL of concentrated RNA (>150 μM) was added to 6 mg 1-ethyl-3-(3-dimethylamino) propyl carbodiimide hydrochloride (EDC), followed by 10 μL of 6 mg/ml Alexa Fluor 488 dissolved in DMSO. Samples were vortexed until the contents completely dissolved before adding 20 μL of 0.1M imidazole pH 5.9. The reactions were incubated at +20°C for 2 hours in the dark, then had the free dye removed using Zeba Spin Desalting Columns (Thermo).

### Microscale thermophoresis

Purified Nsp1 and Nsp1(N127) were serially diluted, with the highest concentration in the assay ranging between 52.1 and 163.2 μM. A constant amount of the fluorescently labelled RNA was added, resulting in a final concentration of 100 nM for the assay. The mixtures were incubated at 20°C for 15 min and loaded into a Monolith NT.155 instrument (Nanotemper Technologies, Munich, Germany) using standard capillaries. Thermophoresis was measured at 25°C and performed using 50% excitation power and medium IR-Laser power. Initial fluorescence was measured from −1.0 to 0 s and used to normalize the measured fluorescent migration time (4.0 to 5.0 s). Three independent replicates were analyzed using MO.Affinity analysis software v2.3 and fit to the standard Kd fit model, which describes a 1:1 stoichiometric molecular interaction according to the law of mass action. The dissociation constant (Kd) is estimated by fitting the equation:

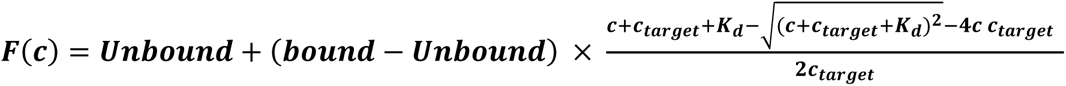

where F(c) is the fraction bound at a given ligand concentration c. Unbound is the Fnorm signal of the isolated target; Bound is the Fnorm signal of the complex, while ctarget is the final concentration of the target in the specific assay.

### Western blotting

Whole-cell lysates were prepared by the direct lysis of PBS-washed cell monolayers with 1x Laemmli sample buffer (50 mM Tris-HCl pH 6.8, 10% glycerol, 2% sodium dodecyl sulphate (SDS), 100 mM DTT, 0.005% Bromophenol Blue). Lysates were immediately placed on ice, homogenized by passing through a 21-gauge needle, and stored at −20°C. Aliquots of lysates thawed on ice were incubated at +95°C for 3 min, cooled on ice, separated using denaturing polyacrylamide gel electrophoresis, transferred onto polyvinylidene fluoride (PVDF) membranes using Trans Blot Turbo Transfer System with RTA Transfer Packs (Bio-Rad Laboratories, Hercules, CA, USA) according to the manufacturer’s protocol, and analyzed by immunoblotting using antibody-specific protocols. Antibodies to the following targets were used: β-actin (1:2000; HRP-conjugated, mouse, Santa Cruz, sc-47778); HA-tag (1:1000; mouse, Cell Signaling, #2367); LARP1 (1:1000; mouse, Santa Cruz, sc-515873). For band visualization, HRP-conjugated horse anti-mouse IgG (Cell Signaling, #7076) was used with Clarity Western ECL Substrate on the ChemiDoc Touch Imaging System (Bio-Rad Laboratories).

### RT-qPCR and 5’RACE

For analysis of reporter RNA levels, total RNA was isolated at 24 h post-transfection using RNeasy Plus Mini Kit (Qiagen Canada, Toronto, ON) according to manufacturer’s protocol. For two-step reverse transcription and qPCR, Luna Script RT SuperMix (NEB Canada, Mississauga, ON) was used to generate cDNA, followed by qPCR using Luna Universal qPCR Master Mix (NEB Canada). The following primers were used: 18S-Left 5’-cgttcttagttggtggagcg; 18S-Rght 5’-ccggacatctaagggcatca; mScarlet-Left 5’-tgatgaacttcgaggacggc; mScarlet-Rght 5’-cgtcaggagggaagttggtg. Relative levels of reporter transcripts (mScarlet) were determined using ΔΔCt method with normalization to 18S rRNA. To verify the precise location of the 5’ ends of reporter RNAs, template switching oligo (A65427, Thermo Fisher Scientific) and the Template Switching RT Enzyme Mix (NEB Canada) were used to perform 5’ rapid amplification of cDNA ends (5’RACE) analysis according to manufacturer protocol. For final product amplification, the following primers were used: RACE-For 5’-aagcagtggtatcaacgcagagt and mRFP-RACE-Rev 5’-gccgtcctcgaagttcatcacg. Amplicons were subject to Sanger sequencing (Azenta, South Plainfield, NJ, USA).

## ACKNOWLEDGEMENTS

T.R.P. acknowledges Canada Research Chair program, NSERC Discovery Grant and Canada Foundation for Innovation programs.

